# Detection and differentiation of *Xanthomonas translucens* pathovars *translucens* and *undulosa* from wheat and barley by duplex quantitative PCR

**DOI:** 10.1101/2023.03.06.531419

**Authors:** Heting Fu, María Constanza Fleitas, Alian Sarkes, Lipu Wang, Yalong Yang, Kher Zahr, Michael W. Harding, David Feindel, Randy Kutcher, Jie Feng

## Abstract

Two probe-based quantitative PCR (qPCR) systems, namely P-Xtt and P-Xtu, were developed for diagnosis of cereal bacterial leaf streak pathogens *Xanthomonas translucens* pv. *translucens* and pv. *undulosa*, respectively. P-Xtt is specific to pv. *translucens*. P-Xtu is specific to pv. *undulosa*, pv. *cerealis*, pv. *secalis* and pv. *pistaciae*. P-Xtt and P-Xtu worked on all accessible strains of pv. *translucens* and pv. *undulosa*, respectively. Both systems could detect 100 copies of the target gBlock DNA. The two systems could be used in both singleplex qPCR and duplex qPCR with similar efficiencies. On genomic DNA from strains of various *X. translucens* pathovars, both singleplex qPCR and duplex qPCR could specifically detect and differentiate pv. *translucens* and pv. *undulosa*. The duplex qPCR could detect pv. *translucens* and pv. *undulosa* from genomic DNA of 1,000 bacterial cells. On infected barley and wheat grain samples, and on one infected wheat leaf sample, the duplex qPCR showed similar efficiency compared to a previously published qPCR system but with the additional capability of pathovar differentiation. The duplex qPCR system developed in this study will be useful in studies on bacterial leaf streak and detection/differentiation of the pathogens.

## Introduction

Bacterial leaf streak (BLS) is a newly re-emerging disease which can cause significant yield loss and reduced grain quality in wheat (*Triticum* spp.) and barley (*Hordeum vulgare* L.). The causal agent is a group of Gram-negative bacteria named *Xanthomonas translucens* infecting a wide range of species of cereal crops, forage and weedy grasses. The disease was first reported on barley in 1917 (Jones et al., 1917) and later on wheat (Smith et al., 1919). In Canada, Drayton (1923) firstly reported the disease on wheat and barley in Manitoba as early as 1922. Over the past two decades, the number of BLS outbreaks has increased dramatically in North America (Curland et al. 2018; Tambong et al. 2021; Arnason 2021).

*Xanthomonas translucens* mainly infects plant species in the Poaceae with the ‘translucens’ group on edible cereal crop species (pathovars: *undulosa*, *translucens*, *cerealis*, *hordei*, and *secalis*) and the ‘graminis’ group (pathovars: *arrhenatheri*, *graminis*, *poae*, *phlei*) on forage grass species (Sapkota et al., 2020). Pathovar (pv.) identification and classification has been based on host range and symptom profiles. Most pathovars have a narrow host range, while a few can infect a wide range of hosts. *Xanthomonas translucens* pv. *undulosa* (*Xtu*) and *X. translucens* pv. *translucens* (*Xtt*) are the most economically important pathovars, with *Xtu* being the most common pathovar affecting wheat whereas *Xtt* is more specific to barley (Sapkota et al. 2020).

Timely and accurate detection and diagnosis of *X. translucens* are critical for the effective management of this pathogen. The pathovar classification of *X. translucens* has presented challenges. While whole genome sequencing and multilocus sequence analysis (Curland et al. 2018; Ledman et al. 2021; Tambong et al. 2021) have been effective in identifying pathovars, they are time-consuming and impractical on plant samples. One loop-mediated isothermal amplification (LAMP) protocol has been developed (Langlois et al. 2017) for the specific detection of three *X. translucens* pathovars, but cannot differentiate among them. Sarkes et al. (2022) developed a protocol for *Xtu* detection from wheat leaf and seed samples by quantitative PCR (qPCR) but the method cannot differentiate *Xtu* from *Xtt* (Sarkes et al., 2022). Two PCR systems, one developed by Román-Reyna et al. (2022) and the other by Alvandi et al. (2023), could specifically detect *Xtt* and *Xtu*, respectively. However, there was no multiplex qPCR system for *Xtt* and *Xtu* detection. Thus, this study was conducted with the objectives to develop a duplex qPCR-based diagnostic protocol that can detect, quantify and differentiate between *Xtt* and *Xtu* and to provide reference data that will allow the protocol to be used immediately by diagnostic labs.

## Materials and Methods

### Chemicals and standard techniques

All chemicals and instrument were purchased from Fisher Scientific Canada (Ottawa, ON) unless otherwise specified. All primers, probes and gBlocks were synthesized by Integrated DNA Technologies (Coralville, IA). PCR was conducted in Promega PCR master mix with a ProFlex PCR system. SYBR Green-based qPCR and probe-based qPCR were conducted in SsoAdvanced universal SYBR Green supermix (Bio-Rad Canada, Mississauga, ON) and PrimeTime gene expression master mix (Integrated DNA Technologies), respectively, in a CFX96 touch real-time PCR detection system (Bio-Rad Canada).

### Bacterial isolates

Isolation of *Xtt* and *Xtu* from wheat and/or barley grains was conducted according to Sarkes et al. (2022). Two *Xtt* strains, namely CDCN-B053 and B054, six *Xtu* strains, namely CDCN-B045, B046, B047, B055, B057 and B058, were isolated from barley and wheat leaf samples. The identities of these isolates were confirmed by sequencing the 16S ribosomal RNA or the *cpn60* gene (Table S1). Among these strains, CDCN-B046 and CDCN-B053 were whole-genome sequenced (GenBank accession numbers JANVBQ010000026 and JANVBP010000007, respectively). PCR amplification of the 16S and *cpn60* followed Klindworth et al. (2013) with the primer pair S-17/A-21 and Sahin et al. (2010) with the primer pair H1594/H1595, respectively. Sequencing of the PCR products was done with a SeqStudio genetic analyzer. Other bacterial species, if not otherwise specified, were retrieved from the plant pathogen culture collection of APHL and their identification was described in Sarkes et al. (2022). All bacterial isolates were maintained on 2.3% (w/v) nutrient agar in darkness at 28°C and as long-term culture in 15% glycerol at -80°C.

### DNA preparation

Two 394-base pair (bp) double-strand DNA fragments, with a sequence of GenBank accession number LT604072 (nt: 4206041 to 4206434) and CP043500 (nt: 4106698 to 4107091), were synthesized as gBlocks and named as G-Xtt and G-Xtu, respectively. Each gBlock was dissolved in water and a stock solution at 8.3 pM was prepared, which roughly equals to 0.5 × 10^7^ molecules/µ L.

All DNA extraction was conducted using the DNeasy Plant Pro kits (Qiagen Canada, Toronto, ON) with a Qiacube (Qiagen Canada). DNA extracted from each sample was eluted in 50 µ L of elution buffer. For DNA extraction from bacteria, a 1-mL cell suspension was centrifuged at 5,000 × *g* for 5 min; the supernatant was removed to leave 100 µ L to avoid disturbing the cell pellet. DNA was extracted from the resultant 100-µ L samples.

Genomic DNA samples of other *X. translucens* strains were kindly provided by researchers in the United States. These included two *Xtt* strains from Dr. Zhaohui Liu, North Dakota State University, Fargo, ND, and four *Xtt* strains, four *Xtu* strains, one pv. *graminis* strain, one pv. *arrhenatheri* strain, one pv. *poae* strain, one pv. *phlei* strain, one pv. *phleipratensis* strain and two pv. *secalis* strains from Rebecca Curland, University of Minnesota, St. Paul, MN. From each DNA sample, a 20-30 µ L subsample was transferred into a new tube and diluted with water to a final volume of 100 µ L. The DNA concentration in each subsample was measured using the NanoDrop 1000 with water as the blanking sample.

### Primer design

In a previous study (Sarkes et al. 2022), 27 genes from the *Xtt* strain DSM18974 were selected as candidates for the development of the qPCR system specific to both *Xtt* and *Xtu*. Among these 27 genes, one gene (GenBank accession number: SCB06379) was found in *Xtt* only, i.e. not present in any other organisms based on the National Center for Biotechnology Information (NCBI) database (https://www.ncbi.nlm.nih.gov) on the time. The genomic sequence of SCB06379 was retrieved and its location on the whole genome sequence of DSM18974 (GenBank accession number: LT604072) was identified. The sequence was extended to its 5’ end and 3’ end, both for 1,000 bp at once, and then the extended sequence was used as the query to search against the *Xtu* (taxid:487909) whole-genome shotgun contigs (wgs) database with the Basic Local Alignment Search Tool (Blast). The sequence extension and Blast were repeated until homologous sequences in the two pathovars were found. By this approach, a heterologous region was identified (Fig. 1). Based on the sequence of SCB06379 and the sequence of an *Xtu* gene located in the heterologous region, two qPCR primer sets were designed, respectively, using Primer-Blast (https://www.ncbi.nlm.nih.gov/tools/primer-blast). For each of the two primer sets, a probe was designed using Primer 3 (https://primer3.ut.ee). The probes in the two primer sets were labelled with different fluorescent dyes (6-FAM for *Xtt* and HEX for *Xtu*). The two primer/probe sets were named P-Xtt and P-Xtu, respectively (Table 1).

**Fig. 1.**
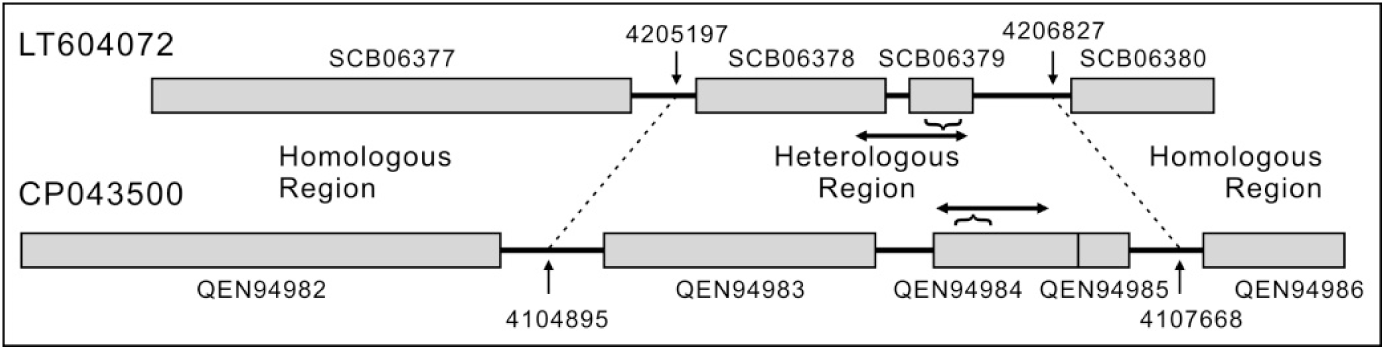
Diagrammatic sketch illustrating the genomic positions of P-Xtt and P-Xtu. The heterologous region, framed by homologous regions, on GenBank accession number LT604072 (*Xanthomonas translucens* pv. *translucens* strain DSM18974) and on GenBank accession number CP043500 (*X. translucens* pv. *undulosa* strain P3) is indicated. The brackets indicate the locations of P-Xtt and P-Xtu. The double-headed arrows indicate the location of the two gBlocks (G-Xtt and G-Xtu).

**Table 1.**
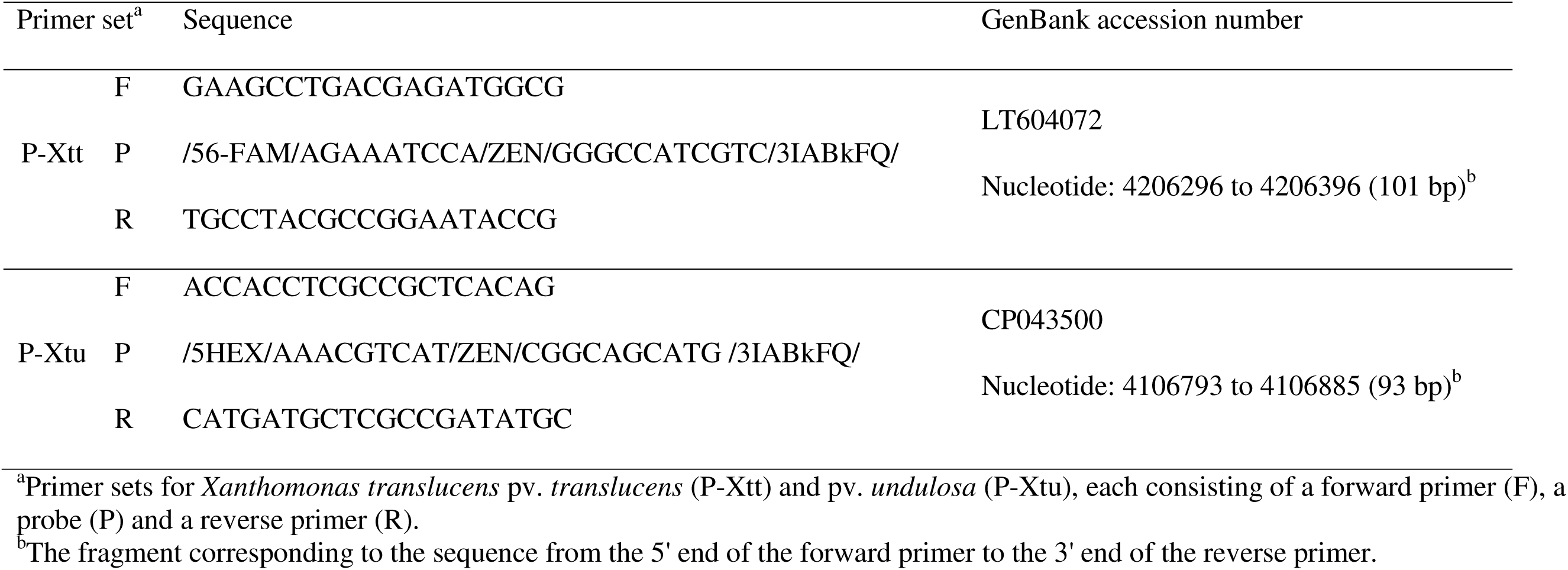
Primers and probes designed in this study

### Primer sequence analysis

All primers and probes were analyzed by Blast against the NCBI nr/nt database. For each primer or probe, the presence of identical sequence (only one alignment, with 100% coverage and 100% identity) in organisms other than *X. translucens* pathovars was checked.

The sequences of the primer target, i.e., the sequence from the 5’ end of the forward primer site to the 3’ end of the reverse primer site, of the two qPCR systems were used as queries to Blast two databases: nr/nt and wgs (Xanthomonas taxid: 338). The Blast results were manually checked for the absence of sequences identical to the queries in species other than *X. translucens* pathovars. In addition, the presence of P-Xtt and P-Xtu in all whole-genome sequenced *Xtt* and *Xtu* was confirmed by checking the presence of all whole-genome sequenced strains in the Blast results. The lists of whole-genome sequenced strains of the two pathovars were obtained by searching “Xanthomonas translucens” in the NCBI genome database (www.ncbi.nlm.nih.gov/genome) and manually checking the resultant Genome Assembly and Annotation reports.

### qPCR reaction

Each qPCR reaction was 20 µ L containing 2 µ L template and 0.25 µ M of each primer. The probe-based qPCR reaction also contained 0.15 µM of each probe. The qPCR program consisted of an initial denaturation step at 95°C for 2 min, followed by 40 cycles of 5 s at 95°C and 30 s at 60°C. Each qPCR reaction was conducted with three technical repeats.

### Specificity test of P-Xtt and P-Xtu on the species level

The specificity of P-Xtt and P-Xtu was tested on templates from the following organisms: *Xtt* strain CDCN-B053, *Xtu* strain CDCN-B046, healthy wheat grains and healthy barley grains, pellets of 20 mL of YPG media (1% yeast extract, 1% peptone, 2% glucose, w/v) after incubating five healthy wheat grains and five healthy barley grains at 150 rpm 30°C for 48 hours (the pellets were prepared by centrifuging the culture after the grains were removed at 5,000 × *g* for 5 min) and eight bacterial cultures on nutrient agar including *Pseudomonas syringae* pv. *atrofaciens*, *P. syringae* pv. *syringae*, *Erwinia rhapontici*, *Clavibacter michiganensis*, *Xanthomonas campestris* pv. *pelargonii*, *Pantoea agglomerans*, *P. allii* and *P. ananatis*. Template DNA in each 20-µ L reaction was 2 ng for bacterial samples or 10 ng for plant samples and the grain culture sample. The experiment was conducted twice with similar results.

### Specificity test of P-Xtt and P-Xtu on the pathovar level

Genomic DNA from ten strains of *Xtu*, eight strains of *Xtt*, and one strain of *X. translucens* pv. *cerealis*, pv. *secalis,* pv. *graminis*, pv. *arrhenatheri*, pv. *poae*, pv. *phlei* and pv. *phleipratensis* were tested by P-Xtu and P-Xtt in both singleplex and duplex qPCR. All templates were also tested with the primer pair specific to *X. translucens* 16S region (F16S/R16S; Sarkes et al. 2022) by SYBR Green qPCR, which served as DNA quantity control. The experiment was conducted twice with similar results.

### Sensitivity test of P-Xtt and P-Xtu on gBlocks

From the stock solution of each of the two gBlocks (G-Xtt and G-Xtu), a set of 10× serial dilutions were prepared from 0.5 × 10^7^ to 5 molecules/µ L (seven solutions for each gBlock). Using the serial dilutions as templates, sensitivity of P-Xtt and P-Xtu was evaluated in both singleplex and duplex qPCR reactions. The experiment was conducted twice with similar results.

### Sensitivity test of P-Xtt and P-Xtu on genomic DNA from bacterial cell dilutions

Cell suspension was prepared from the *Xtt* strain CDCN-B053 and the *Xtu* strain CDCN-B046. The suspensions were adjusted with water to OD600 = 0.6, which was equivalent to 1 × 10^9^ cells/mL, as confirmed by counting of the 100× and 1,000× dilutions using a haemocytometer. From the 1 × 10^9^ cells/mL mixture a set of 10× cell serial dilutions were prepared until 1 × 10^3^ cells/mL. DNA was extracted from three 1-mL samples (three biological repeats) of each dilution. All DNA samples were tested by duplex qPCR using P-Xtt/P-Xtu and singleplex qPCR using the F11/R11/P11 primers/probe set developed by Sarkes et al. (2022). The experiment was conducted twice with similar results.

### Detection of *Xtt* and *Xtu* from plant samples

Six barley and five wheat samples were tested by duplex qPCR using P-Xtt/P-Xtu and singleplex qPCR using the F11/R11/P11 primers/probe set. Preparation of grain samples for DNA extraction followed Sarkes et al. (2022). Briefly, 100 kernels from each sample were soaked in 4 mL water in a 50-mL tube. The tube was vortexed for 30 sec, from which a 1-mL sample of supernatant was transferred to a 1.5-mL tube. The tube was centrifuged at 11,600 × *g* for 2 min and 900 µ L of the supernatant removed. DNA was extracted from the remaining 100 µ L pellet/supernatant mixture, which represented all extractible DNA from the surface of 25 kernels. For DNA extraction from the wheat leaf sample, a 2-cm fragment of necrotic leaf was used.

### Data analysis

In the test on DNA from the bacterial cell dilutions, means of the quantification cycle (Cq) values were calculated from the three technical repeats of each DNA samples. The average of the Cq means from the three biological repeats of each cell dilution was regarded as one data point. For all other qPCR, the Cq mean from the three technical repeats was regarded as one data point. The qPCR standard curves were constructed by regression analysis using the SAS software (version 9.4; SAS Institute, Cary, NC). The qPCR primer efficiencies were calculated as E = - 1+10^(-1/slope)^ (Svec et al. 2015). For the same DNA template, the Cq values of singleplex qPCR and duplex qPCR were compared by Student’s *t*-test using the SAS software.

## Results

### Sequence analysis of the specificity of P-Xtt and P-Xtu

Blast of the primer and probe sequence against NCBI nr/nt database indicated that the forward primer and the reverse primer of P-Xtt were present in one non-*X. translucens* organism, respectively, and that the forward primer and the probe of P-Xtu were present in two and four non-*X. translucens* organisms, respectively. However, all of these non-*X. translucens* were different species. The probe of P-Xtt and the reverse primer of P-Xtu were present exclusively in *X. translucens* pathovars. The results also indicated the specificity of P-Xtt and P-Xtu at the pathovar levels, which were identical with the results from Blast of the primer targets of P-Xtt and P-Xtu as described below. The above results were confirmed on May 9, 2023.

The primer targets of P-Xtt and P-Xtu are 101 and 93 bp, respectively (Table 1). Using the two fragments as queries, Blast against NCBI nr/nt database and wgs (Xanthomonas taxid: 338) database was conducted (the results were verified on May 9, 2023; Table S2). For both fragments, Blast against the two databases covered all whole-genome sequenced *Xtt* and *Xtu*, and did not generate any hit from a non-*X. translucens* organism (Table S2). Orthologs with identical sequences as the P-Xtt target were found in 23 out of the 27 whole-genome sequenced *Xtt* strains, one strain of *X. translucens* pv. *hordei* and seven strains of unknown *X. translucens* pathovars. Orthologs were also found in nine strains of pv. *graminis* and three strains of pv. *arrhenatheri*, with identical sequences to each other but were different with the P-Xtt target on six nucleotides (nt). Among the six nt, three were located on the site of the forward primer target (with a sequence of GAAACCTGCCGAAATGGCC) and the other three were located on non-primer/probe-target regions. The P-Xtu target was found in all the whole-genome sequenced strains of *Xtu* (18), *X. translucens* pv. *cerealis* (3), pv. *secalis* (2) and pv. *pistaciae* (1), as well as 17 strains of unknown *X. translucens* pathovars. Orthologs of the P-Xtu target are also present in the four *Xtt* strains that lack the P-Xtt target: strains XtKm33, BLSB3, SIMT-07 and SLV-2. These four strains are *Xtu* but were mislabeled as *Xtt* in the NCBI database. The P-Xtu target and all orthologs had identical sequences.

P-Xtt and P-Xtu, as well as their targets, are not present in whole-genome sequenced *X. translucens* pv. *poae*, pv. *phlei* or pv. *phleipratensis.* These results indicated that P-Xtt is specific to *Xtt* and and *X. translucens* pv. *hordei*, P-Xtu is specific to *Xtu*, *X. translucens* pv. *cerealis*, pv. *secalis* and pv. *pistaciae*, and both P-Xtt and P-Xtu are ubiquitous among strains of the specific pathovars. It has not escaped our notice that the whole genome sequence of *X. translucens* pv. *hordei* strain UPB458 was relabeled as *Xtt* (Table S1). Thus, based on the database-available genomic sequences, we can conclude that P-Xtt is specific to *Xtt* only.

### Specificity of P-Xtt and P-Xtu at the species level

When P-Xtt and P-Xtu were tested on DNA from healthy (non-symptomatic) wheat and barley, microorganisms associated with the kernel surface, and eight other bacterial species, no positive signal was observed (Table 2). This result indicated that both P-Xtt and P-Xtu are specific to *X. translucens* and DNA from host plants or other bacteria commonly associated with wheat or barley kernels will not interfere with the qPCR reactions.

**Table 2.** Specificity test of P-Xtt and P-Xtu on DNA of *Xanthomonas translucens* pv. *translucens* and pv. *undulosa* and other organisms

| DNA template<sup>a</sup> | Cq value (mean ± standard deviation; n = 3) |  |
| --- | --- | --- |
|  | P-Xtt | P-Xtu |
| CDCN-B046 (*X. translucens* pv. *undulosa*) | ns<sup>c</sup> | 24.9±0.1 |
| CDCN-B053 (*X. translucens* pv. *translucens*) | 23.6±0.1 | ns |
| Healthy wheat leaves | ns | ns |
| Healthy barley leaves | ns | ns |
| Healthy wheat grains | ns | ns |
| Health barley grains | ns | ns |
| Grain culture in YPG medium<sup>b</sup> | ns | ns |
| *Pseudomonas syringae* pv. *atrofaciens* | ns | ns |
| *P. syringae* pv. *syringae* | 38.7<sup>d</sup> | ns |
| *Erwinia rhapontici* | ns | ns |
| *Clavibacter michiganensis* | ns | ns |
| *Xanthomonas campestris* pv. *pelargonii* | 39.3<sup>d</sup> | ns |
| *Pantoea agglomerans* | ns | ns |
| *P. allii* | ns | ns |
| *P. ananatis* | ns | ns |

<sup>a</sup>DNA template in each 20-μL qPCR reaction: bacteria, 2 ng; plants and the grain culture, 10 ng.
<sup>b</sup>DNA from microorganisms grown in YPG media (1% yeast extract, 1% peptone, 2% glucose, w/v) after incubating healthy wheat and barley grains at 150 rpm 30°C for 48 hours.
<sup>c</sup>ns = no signal from all of the three technical repeats.
<sup>d</sup>Only one of the three technical repeats produced a Cq value.

### Specificity of P-Xtt and P-Xtu on bacterial DNA

DNA from strains of *X. translucens* pathovars were tested by P-Xtt and P-Xtu by singleplex and duplex qPCR (Table 3). Positive signals (mean Cq value <25) were produced by P-Xtt from all *Xtt* strains but not from other pathovars. Cq values >37 were produced from some strains of other pathovars, however, in most cases only from one or two of the three technical repeats. Similarly, positive signals (mean Cq value <28) were produced by P-Xtu from all *Xtu*, *X. translucens* pv. *cerealis* and pv. *secalis* strains but not from other pathovars. Cq values >37 were produced from some strains of other pathovars, however, in most cases only from one or two of the three technical repeats. Both P-Xtt and P-Xtu did not produce positive signals from *X. translucens* pv. *poae*, pv. *phlei* and pv. *phleipratensis*. These results confirmed that P-Xtt is specific to *Xtt*, P-Xtu is specific to *Xtu*, *X. translucens* pv. *cerealis* and pv. *secalis*, and both P-Xtt and P-Xtu are ubiquitous among strains of the specific pathovars.

**Table 3.**
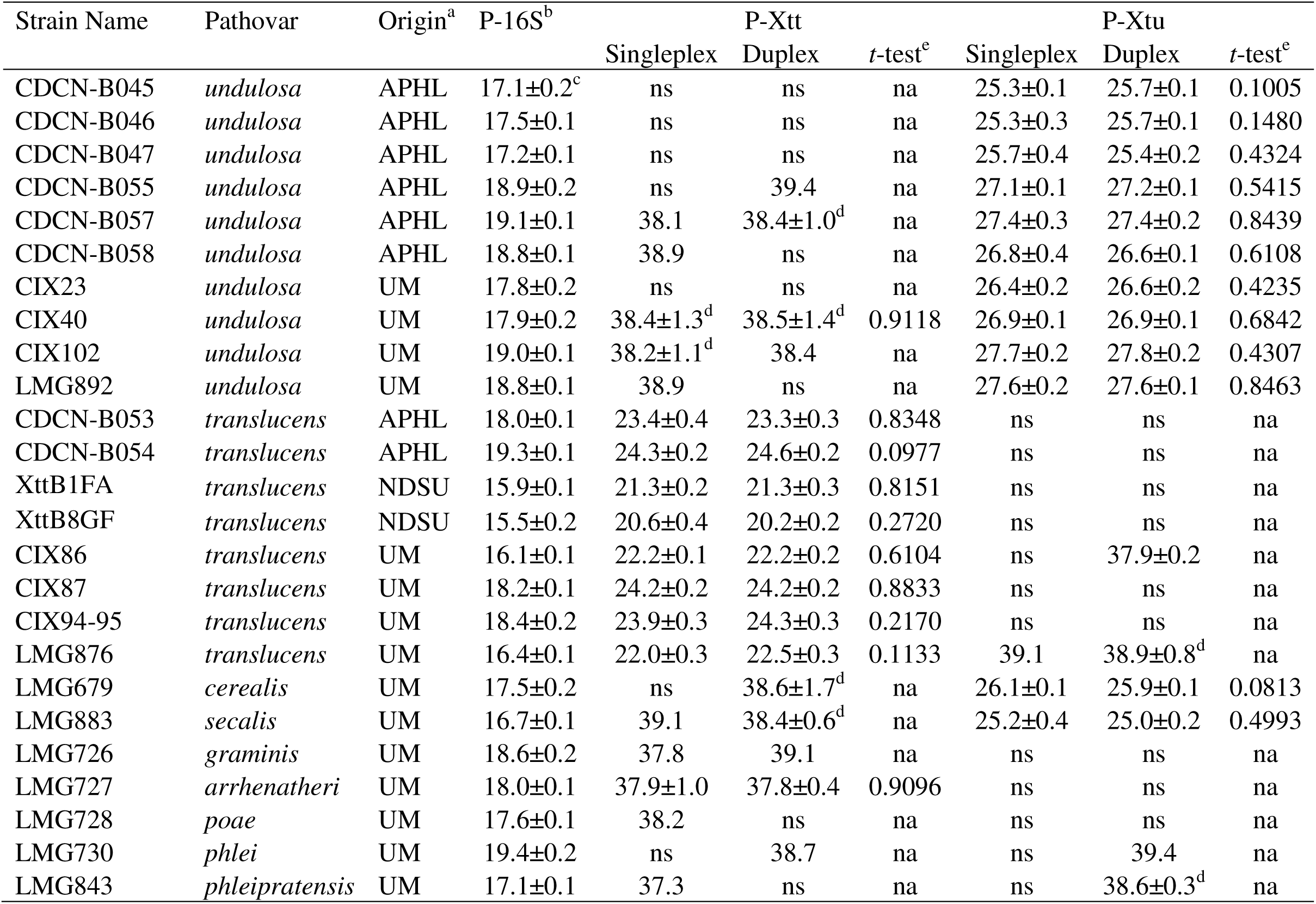
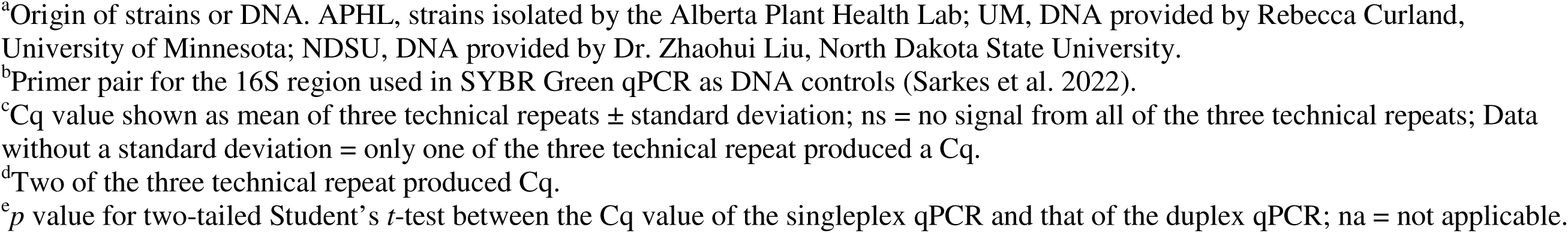
Specificity test of P-Xtt and P-Xtu on DNA of *Xanthomonas translucens* pathovars by singleplex and duplex qPCR

| Strain Name | Pathovar | Origin<sup>a</sup> | P-16S<sup>b</sup> | P-Xtt |  |  | P-Xtu |  |  |
| --- | --- | --- | --- | --- | --- | --- | --- | --- | --- |
|  |  |  |  | Singleplex | Duplex | *t*-test<sup>c</sup> | Singleplex | Duplex | *t*-test<sup>c</sup> |
| CDCN-B045 | *undulosa* | APHL | 17.1±0.2<sup>c</sup> | ns | ns | na | 25.3±0.1 | 25.7±0.1 | 0.1005 |
| CDCN-B046 | *undulosa* | APHL | 17.5±0.1 | ns | ns | na | 25.3±0.3 | 25.7±0.1 | 0.1480 |
| CDCN-B047 | *undulosa* | APHL | 17.2±0.1 | ns | ns | na | 25.7±0.4 | 25.4±0.2 | 0.4324 |
| CDCN-B055 | *undulosa* | APHL | 18.9±0.2 | ns | 39.4 | na | 27.1±0.1 | 27.2±0.1 | 0.5415 |
| CDCN-B057 | *undulosa* | APHL | 19.1±0.1 | 38.1 | 38.4±1.0<sup>d</sup> | na | 27.4±0.3 | 27.4±0.2 | 0.8439 |
| CDCN-B058 | *undulosa* | APHL | 18.8±0.1 | 38.9 | ns | na | 26.8±0.4 | 26.6±0.1 | 0.6108 |
| CIX23 | *undulosa* | UM | 17.8±0.2 | ns | ns | na | 26.4±0.2 | 26.6±0.2 | 0.4235 |
| CIX40 | *undulosa* | UM | 17.9±0.2 | 38.4±1.3<sup>d</sup> | 38.5±1.4<sup>d</sup> | 0.9118 | 26.9±0.1 | 26.9±0.1 | 0.6842 |
| CIX102 | *undulosa* | UM | 19.0±0.1 | 38.2±1.1<sup>d</sup> | 38.4 | na | 27.7±0.2 | 27.8±0.2 | 0.4307 |
| LMG892 | *undulosa* | UM | 18.8±0.1 | 38.9 | ns | na | 27.6±0.2 | 27.6±0.1 | 0.8463 |
| CDCN-B053 | *translucens* | APHL | 18.0±0.1 | 23.4±0.4 | 23.3±0.3 | 0.8348 | ns | ns | na |
| CDCN-B054 | *translucens* | APHL | 19.3±0.1 | 24.3±0.2 | 24.6±0.2 | 0.0977 | ns | ns | na |
| XttB1FA | *translucens* | NDSU | 15.9±0.1 | 21.3±0.2 | 21.3±0.3 | 0.8151 | ns | ns | na |
| XttB8GF | *translucens* | NDSU | 15.5±0.2 | 20.6±0.4 | 20.2±0.2 | 0.2720 | ns | ns | na |
| CIX86 | *translucens* | UM | 16.1±0.1 | 22.2±0.1 | 22.2±0.2 | 0.6104 | ns | 37.9±0.2 | na |
| CIX87 | *translucens* | UM | 18.2±0.1 | 24.2±0.2 | 24.2±0.2 | 0.8833 | ns | ns | na |
| CIX94-95 | *translucens* | UM | 18.4±0.2 | 23.9±0.3 | 24.3±0.3 | 0.2170 | ns | ns | na |
| LMG876 | *translucens* | UM | 16.4±0.1 | 22.0±0.3 | 22.5±0.3 | 0.1133 | 39.1 | 38.9±0.8<sup>d</sup> | na |
| LMG679 | *cerealis* | UM | 17.5±0.2 | ns | 38.6±1.7<sup>d</sup> | na | 26.1±0.1 | 25.9±0.1 | 0.0813 |
| LMG883 | *secalis* | UM | 16.7±0.1 | 39.1 | 38.4±0.6<sup>d</sup> | na | 25.2±0.4 | 25.0±0.2 | 0.4993 |
| LMG726 | *graminis* | UM | 18.6±0.2 | 37.8 | 39.1 | na | ns | ns | na |
| LMG727 | *arrhenatheri* | UM | 18.0±0.1 | 37.9±1.0 | 37.8±0.4 | 0.9096 | ns | ns | na |
| LMG728 | *poae* | UM | 17.6±0.1 | 38.2 | ns | na | ns | ns | na |
| LMG730 | *phlei* | UM | 19.4±0.2 | ns | 38.7 | na | ns | 39.4 | na |
| LMG843 | *phleipratensis* | UM | 17.1±0.1 | 37.3 | ns | na | ns | 38.6±0.3<sup>d</sup> | na |

<sup>a</sup>Origin of strains or DNA. APHL, strains isolated by the Alberta Plant Health Lab; UM, DNA provided by Rebecca Curland, University of Minnesota; NDSU, DNA provided by Dr. Zhaohui Liu, North Dakota State University.
<sup>b</sup>Primer pair for the 16S region used in SYBR Green qPCR as DNA controls (Sarkes et al. 2022).
<sup>c</sup>Cq value shown as mean of three technical repeats ± standard deviation; ns = no signal from all of the three technical repeats; Data without a standard deviation = only one of the three technical repeat produced a Cq.
<sup>d</sup>Two of the three technical repeat produced Cq.
<sup>e</sup>*p* value for two-tailed Student's *t*-test between the Cq value of the singleplex qPCR and that of the duplex qPCR; na = not applicable.

Cq values generated by singleplex qPCR and duplex qPCR were compared by Student’s *t*-test. Among all the applicable samples, *p* values between 0.0977 and 0.9118 were obtained. These results indicated that duplex qPCR using P-Xtt and P-Xtu in the same reaction had the same robustness as singleplex qPCR on detection and differentiation of *Xtt* and *Xtu*.

### Sensitivity of P-Xtt and P-Xtu on gBlocks

The serial dilutions of the gBlock G-Xtt were tested in singleplex qPCR using P-Xtt and in duplex qPCR using P-Xtt/P-Xtu. Similarly, G-Xtu was tested with P-Xtu and P-Xtt/P-Xtu. Based on the Cq values, a standard curve was constructed for each primer set (Fig. 2). The efficiencies of P-Xtt in both singleplex and duplex qPCR were 0.98, respectively (Fig. 2a and 2b). The efficiencies of P-Xtu in singleplex and duplex qPCR were 1.04 (Fig. 2c) and 1.03 (Fig. 2d), respectively. For P-Xtt, on each concentration of the gBlock serial dilutions, the Cq value from singleplex qPCR (Fig. 2a) and that from duplex qPCR (Fig. 2b) were compared by two-tailed Student’s *t*-test. Of all gBlock concentrations, the *p* values of the *t*-test were in the range of 0.1233 to 0.9095, indicating no statistically significant difference in Cq between singleplex and multiplex reactions. Similar results were observed for P-Xtu (Fig. 2c and d), in which the *p* values were in the range of 0.2416 to 0.9974. The results confirmed that P-Xtt and P-Xtu can be used in duplex qPCR and singleplex qPCR with the same robustness.

**Fig. 2.**
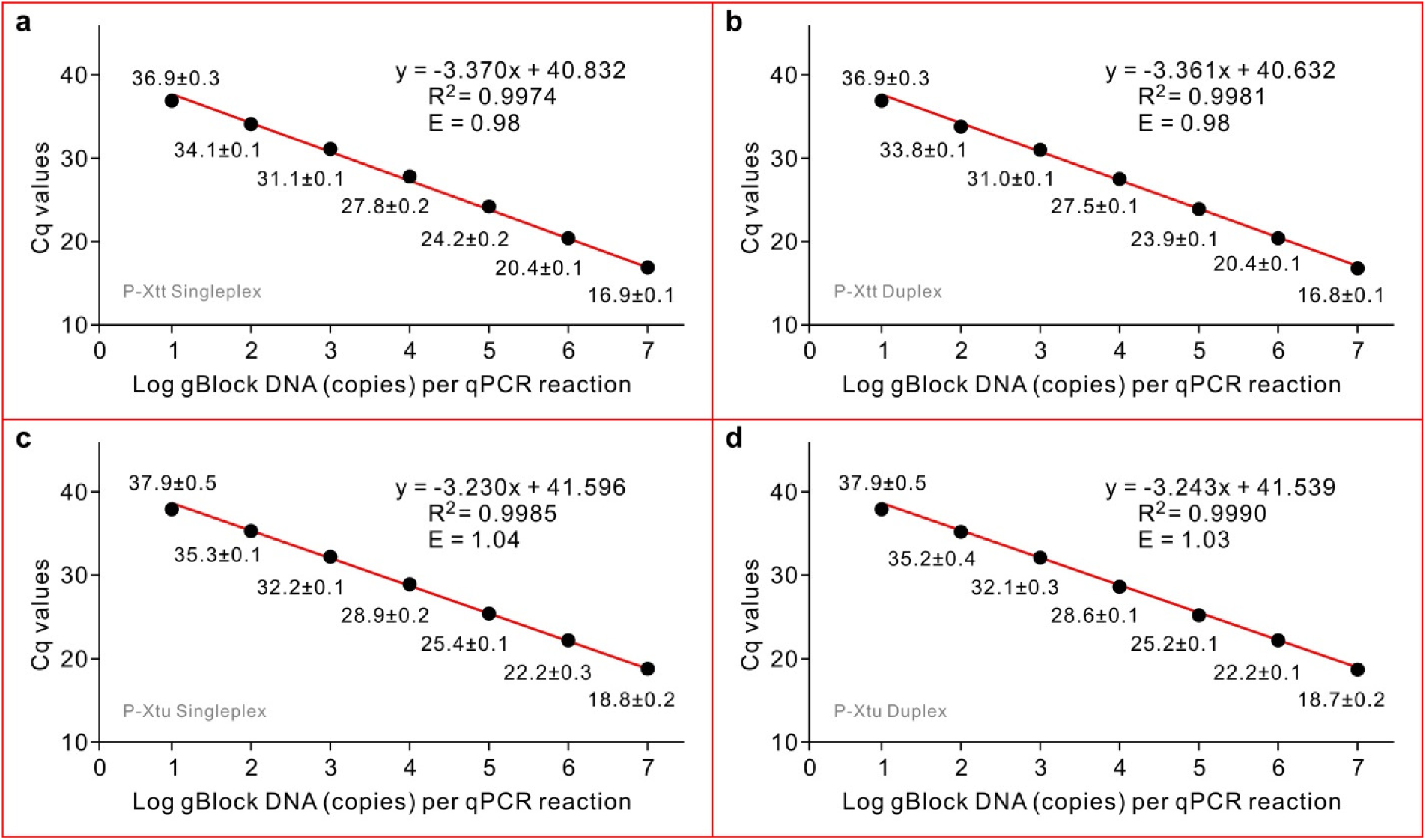
Sensitivity of the qPCR systems P-Xtt for *Xanthomonas translucens* pv. *translucens* and P-Xtu for pv. *undulosa* on gBlock DNA. a and b, P-Xtt singleplex qPCR and P-Xtt/P-Xtu duplex qPCR on the pv. *translucens* gBlock, respectively. c and d, P-Xtu singleplex qPCR and P-Xtt/P-Xtu duplex qPCR on the pv. *undulosa* gBlock, respectively. The qPCR standard curves were generated from the mean of quantification cycle (Cq) values against log10 of gBlock DNA copies in one reaction. The R^2^ score of the equation and the efficiency of the primers (E) are indicated over the curve. Efficiency was calculated as E = -1+10^(-1/slope)^. Each data point is shown as mean of three technical repeats ± standard deviation.

### Sensitivity of P-Xtt and P-Xtu on DNA extracted from bacterial cell dilutions

When DNA from bacterial cell dilutions was tested by the duplex qPCR, the lowest number of cells that could be detected by one reaction was 10^2.6^ for both *Xtt* and *Xtu* (Fig. 3). On three DNA samples (biological repeats) each extracted from 10^2.6^ cells, P-Xtt and P-Xtu generated Cq values of 36.3 and 39.1, respectively. The efficiencies of both P-Xtt and P-Xtu were 0.79 (Fig. 3).

**Fig. 3.**
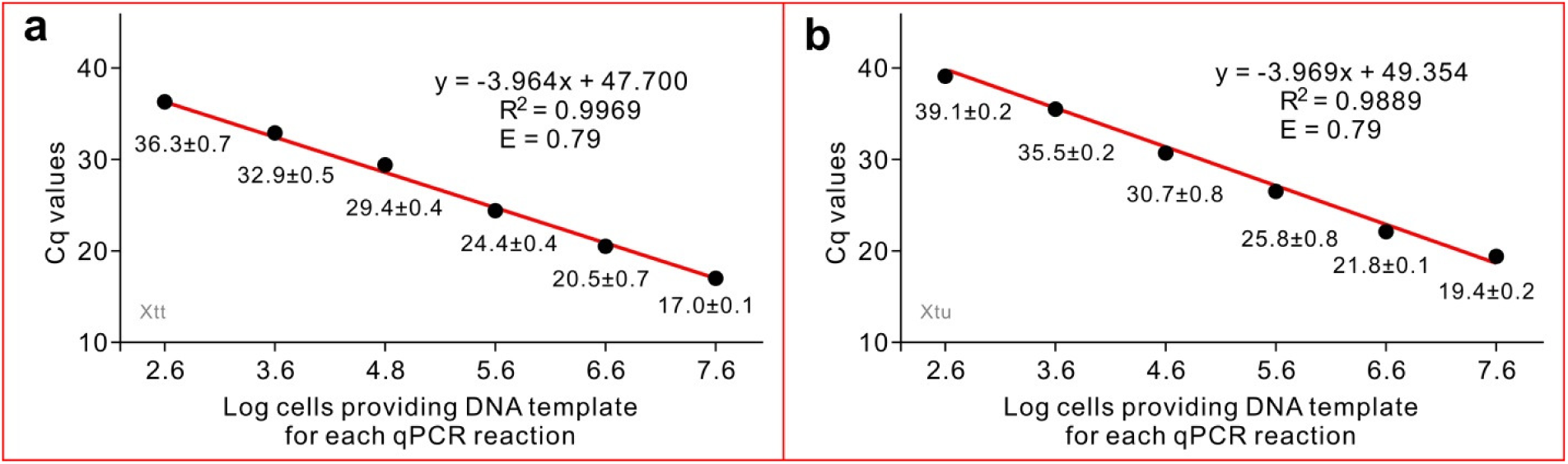
Sensitivity of the duplex qPCR system on the detection of *Xanthomonas translucens* pv. *translucens* (a) and pv. *undulosa* (b). The qPCR standard curves were generated from the mean of quantification cycle (Cq) values against log10 of the cell number from which the DNA was used in one reaction. The R^2^ score of the equation and the efficiency of the primers (E) are indicated over the curve. Efficiency was calculated as E = -1+10^(-1/slope)^. Each data point is shown as mean of three biological repeats ± standard deviation.

### Detection of *Xtt* and *Xtu* from plant samples

Both duplex qPCR using P-Xtt/P-Xtu and singleplex qPCR using F11/P11/R11 detected *Xtt* from the six barley samples and *Xtu* from the five wheat samples (Table 4). From the barley sample B4, *Xtu* was also detected by the duplex qPCR. This result was supported by bacterial isolation: both *Xtt* and *Xtu* were isolated from this sample and the identities of the isolates were confirmed by the sequences of the *cpn60* gene.

**Table 4.** Test of barley and wheat samples by duplex qPCR

| Sample<sup>a</sup> | Type | Origin<sup>b</sup> | Primers<sup>c</sup> |  |  |
| --- | --- | --- | --- | --- | --- |
|  |  |  | P-Xtt | P-Xtu | F11/P11/R11 |
| B1 | Grain | Sturgeon | 21.2±0.2 | 39.7±0.2 | 22.1±0.2 |
| B2 | Grain | Lacombe | 25.6±0.1<sup>d</sup> | No signal | 27.2±0.1 |
| B3 | Grain | Lacombe | 21.2±0.1 | No signal | 22.3±0.2 |
| B4 | Grain | Lethbridge | 25.6±0.1 | 25.9±0.1 | 25.0±0.1 |
| B5 | Grain | Lethbridge | 20.7±0.3 | 39.0<sup>e</sup> | 21.1±0.1 |
| B6 | Grain | Taber | 21.6±0.2 | No signal | 22.5±0.1 |
| W1 | Leaf | USASK | No signal | 22.3±0.2 | 20.3±0.1 |
| W2 | Grain | Lethbridge | No signal | 28.0±0.1 | 25.8±0.1 |
| W3 | Grain | Forty Mile | No signal | 22.3±0.1 | 20.2±0.1 |
| W4 | Grain | Leduc | No signal | 26.6±0.2 | 28.1±0.1 |
| W5 | Grain | Table | 39.4<sup>e</sup> | 31.9±0.1 | 29.9±0.3 |
| B053 | Culture | CDCN | 23.3±0.2 | No signal | 22.7±0.2 |
| B046 | Culture | CDCN | No signal | 26.6±0.1 | 24.5±0.2 |

<sup>a</sup>Six barley grain samples (B1 to B6), one infected wheat leaf sample (W1), four wheat grain samples (W2 to W5), and the culture of two whole-genome sequenced *X. translucens* strains: pv. *translucens* CDCN-B053 and pv. *undulosa* CDCN-B046.
<sup>b</sup>Origin of the samples indicated as counties of Alberta, University of Saskatchewan (USASK) or Crop Diversification Centre North (CDCN).
<sup>c</sup>P-Xtt and P-Xtu were used in duplex qPCR; the primers/probe set F11/P11/R11 (Sarkes et al. 2022) was used as control.
<sup>d</sup>Cq values shown as mean of three technical repeats ± standard deviation.
<sup>e</sup>Only one of the three technical repeats produced a Cq.

## Discussion

### Specificity of P-Xtt and P-Xtu

The target of P-Xtt was originally identified by Sarkes et al. (2022). It was then present in all whole genome sequenced *Xtt* strains and not in other *X. translucens* pathovars. After the release of the whole genome sequences of *X. translucens* pv. *arrhenatheri* and pv. *graminis,* orthologs of the P-Xtt target were found in these genomes. The sequences of these orthologs are identical to each other but 6-nt different to the P-Xtt target. These orthologs carry the sequences of the reverse primer and the probe of P-Xtt, but have 3-nt difference to the forward primer on the forward primer site, with one of them at the 3’ end. On the DNA of a pv. *arrhenatheri* strain and a pv. *graminis* strain, P-Xtt did not generate positive signal (Table 3), indicating that the 3-nt difference on the forward primer is sufficient to ensure the specificity of P-Xtt to *Xtt*. Thus, based on the current database, P-Xtt is specific to *Xtt* at the pathovar level and ubiquitous at the strain level. A qPCR system 62xtt558F/62xtt616R/62xtt579pb specific to *Xtt* was recently developed (Tambong et al. 2023). However, the 78-nt target of this system is not present in some of the whole genome sequenced *Xtt* strains (e.g. strains AB16-8-8a, BC13-4-5a, etc.). In addition, polymorphisms are present at the reverse primer (1 nt) and the probe (2 nt) sites in some *Xtt* strains (e.g. strains CDCN-B053, etc.). Whether these polymorphisms will compromise the qPCR efficiency is unknown, however, similar polymorphisms are also present in all whole genome sequenced pv. *poae* strains.

Orthologs identical to the P-Xtu target are present in all whole genome sequenced strains of *Xtu*, *X. translucens* pv. *secalis*, pv. *cerealis* and pv. *pistaciae* (Table S2). None of the previously published *Xtu*-specific PCR primers are exclusively specific to *Xtu*. For example, the target of the S8.pep-F/S8.pep-R primers (Román-Reyna et al. 2022) is also present in one (strain CFBP2539) of the two whole genome sequenced pv. *secalis* strains with 100% identity. In addition, orthologs are present in all whole genome sequenced strains of pv. *cerealis* and pv. *pistaciae*, albeit there is 1-nt difference in the middle of the forward primer. The conventional PCR system XtuF/XtuR developed by Alvandi et al. (2023) is the most *Xtu*-specific system so far. The sequences of the target are present in all whole genome sequenced *Xtu* strains but not in other organisms except the *X. translucens* pv. *secalis* strain CFBP 2539. Based on the target sequences, we designed a set of qPCR primers/probe. However, compared to singleplex qPCR, compromised results were obtained when this primers/probe set was used with P-Xtt in duplex qPCR. Alternative primers/probes will be designed and tested in future studies.

There are four strains (strains XtKm33, BLSB3, SIMT-07 and SLV-2) labeled as *Xtt* in the NCBI database that carry the P-Xtu target but not the P-Xtt target. These four strains had the same genotype as *Xtu* rather than *Xtt* in the target of the qPCR system F11/R11/P11 developed by Sarkes et al. (Table S3 in Sarkes et al. 2022). The targets of the *Xtu*-specific primer systems S8.pep-F/S8.pep-R primers (Román-Reyna et al. 2022) and XtuF/XtuR (Alvandi et al. 2023) are also present in these four strains. In addition, phylogenetic studies (Khojasteh et al. 2019; Shah et al. 2021) indicated that these four strains are closely related to *Xtu* rather than *Xtt*. These observations lead to a suggested conclusion that these four strains are *Xtu* but mislabeled as *Xtt* in the NCBI database.

Among all the tested organisms (Tables 2 and 3), P-Xtt was specific to *Xtt* and P-Xtu was specific to *Xtu*, *X. translucens* pv. *secalis* and pv. *cerealis*, which was in agreement with the sequence analyses. According to Sapkota et al (2020), *Xtt* is pathogenic to barley, *Xtu* is pathogenic to wheat and barley, *X. translucens* pv. *secalis* is pathogenic to rye and pv. *cerealis* is pathogenic to wheat, barley, oat and brome grass. *Xanthomonas translucens* pv. *pistaciae* is pathogenic to pistachio only (Giblot-Ducray et al. 2009). Thus, in our study, while the positive signal generated by P-Xtt indicated the presence of *Xtt*, the positive signal generated by P-Xtu from barley or wheat samples might likely indicate the presence of *Xtu* or/and *X. translucens* pv. *cerealis*. The PCR target of the primer pair XtuF/XtuR (Alvandi et al. 2023) was not present in any of the pv. *cerealis* genomes. Thus XtuF/XtuR can be used to confirm the identity of the pathovar in the barley sample from which P-Xtu generated positive signal.

### Sensitivities of P-Xtt and P-Xtu

On gBlock DNA, the low limit of both P-Xtt and P-Xtu for a positive detection in both singleplex and duplex qPCR was 10 double stranded DNA copies per reaction (Fig. 2). However, with consideration of the qPCR results from bacterial genomic DNA in which a Cq value of 37 could be produced from non-targeted strains (Table 3), we concluded that the duplex qPCR system can reliably detect 100 double stranded DNA copies. The number of bacterial cells that could ensure the obtaining of 100 copies of genomic DNA must be larger than 100, e.g., approximately 1,000 as indicated in this study (Fig. 3). In a previous study, DNA extracted from 40 bacterial cells could be detected by the qPCR system F11/R11/P11 (Sarkes et al. 2022). The difference on the detection limits of the current study and Sarkes et al. (2022) can be explained by two reasons. First, the robustness of P-Xtt and P-Xtu, in singleplex or in duplex qPCR, might be lower than that of F11/R11/P11, as indicated by the standard curves against gBlock DNA. Second, the efficiencies of DNA extraction in the two studies were different, as indicated by the standard curves against genomic DNA from cell dilutions (Fig. 3 in the current study and Fig. 3c in Sarkes et al. 2022). The possible factors affecting DNA extraction efficiency could be alternative batches of Plant Pro kits and alternative models of Qiacube used in the two studies. In addition to the above two reasons, alternative batches and the storing time of the qPCR master mix might also influence the qPCR efficiency.

After we noticed the difference between the results from the current study and those from Sarkes et al. (2022), regarding the detection limit of cell numbers, tests of the duplex qPCR system on cell dilutions and field samples were repeated with the F11/R11/P11 system being included as control. Against the same template of genomic DNA, the Cq values of F11/R11/P11 (Fig. S1) were lower than those of P-Xtt (in a range of 0.2-1.3; Fig 3a) and P-Xtu (in a range of 0.5-2.5; Fig 3b). Against the same templates of field samples, the Cq values of F11/R11/P11 were either higher or lower than those of P-Xtu (in a range of 1.5 to 2.2). In contrast, the Cq values of F11/R11/P11 were higher than those of P-Xtt (in a range of 0.4-1.2), except the sample B4 from which both *Xtt* and *Xtu* were identified. However, against genomic DNA from the *Xtt* strain CDCN-B053, the Cq value of F11/R11/P11 was lower than that of P-Xtt. These observations might be due to the combined effect of differences in primers’ efficiencies, and different interactions of P-Xtt, P-Xtu and F11/R11/P11 to the background DNA in the tested samples. Perhaps, the overall conclusion on the sensitivity of the duplex qPCR system could be that it is similar or lower than the sensitivity of F11/R11/P11 and with the consideration of DNA extraction efficiencies, the low limit of a positive detection and differentiation of *Xtt* and *Xtu* is 1,000 cells per reaction.

The sensitivity of a PCR-based diagnostic protocol should include not only the efficiency of the primers/probe, but also the sample preparation and the efficiency of DNA extraction. The influence of DNA extraction on the efficiency of qPCR detection systems was previously studied (Yang et al. 2021) and illustrated in the current study. In the current study, we used gBlock DNA to evaluate the primer efficiencies and DNA from cell dilutions to evaluate the efficiency of the entire qPCR system. Many previous studies used serial dilutions from one DNA preparation, rather than DNA samples from serial dilutions of cells, to generate the qPCR standard curve and the resultant data was used to represent both primer efficiency and the efficiency of the qPCR system. We have found that the reported detection limits of some qPCR systems targeting single-copied DNA fragments were lower than the molecular weight of the genome of the target species. This was because of inaccurate concentration measurement of the original DNA sample from which the serial dilutions were prepared and the fact that diluting of a DNA sample also dilutes the PCR inhibitors inherited from the DNA extraction, which will increase the calculated primer efficiency. Therefore, we preferred to use DNA from cell dilutions to evaluate the overall efficiency of the qPCR system, which is more representative of DNA obtained from diagnostic samples compared to DNA dilution series originated from one DNA preparation.

### The usefulness of the duplex qPCR

The P-Xtt/P-Xtu system is the first duplex qPCR system for *Xtt* and *Xtu* detection and differentiation. Compared to other systems, the most important advantage of the duplex qPCR is the capability of quantification of both *Xtt* and *Xtu* simultaneously from one DNA sample. The system will be extremely useful in studies on the population dynamics of *Xtt* and *Xtu*, as well as screening of genetic resources for BLS resistance. Such studies generally include artificial inoculation of *Xtt* and/or *Xtu* in greenhouses or small field trials, thus the fact that P-Xtu is specific to not only *Xtu* but also *X. translucens* pv. *cerealis* and pv. *secalis* will not be a concern. The system is also preferable by diagnostic labs: three major plant diagnostic labs in Alberta, Canada have adopted this system for BLS testing. *Xanthomonas translucens* pv. *cerealis* is pathogenic to both barley and wheat, and pv. *secalis* is phylogenetically clustered with *Xtu* (Sapkota et al. 2020). Thus, if P-Xtu generates positive signal from a wheat sample, no matter the pathogen is *Xtu*, *X. translucens* pv. *cerealis* or pv. *secalis*, attention should be paid on the field or seed storage from which the sample was derived.

According to Sapkota et al. (2020), *Xtu* is pathogenic to both wheat and barley. In the current study, *Xtu* was detected and isolated from one barley sample (B4 in Table 4). The identity of the isolated *Xtu* was confirmed by the *cpn60* sequence, which was at 454-bp, one bp different to the *cpn60* of CDCN-B055 and identical to the *cpn60* of all other *Xtu* isolates listed in Table S1. However, whether the detected *Xtu* was actually pathogenic on that barley sample was not known. So far there is no study indicating whether *Xtt* is also pathogenic to wheat. However, it is possible to find *Xtt* on wheat samples if the wheat field where the samples were derived is adjacent to an *Xtt-*infected barley field. Our duplex qPCR system provides a convenient method to monitor the occurrence of both pathovars. The prevalence of *Xtt* on wheat or *Xtu* on barley will provide useful information guiding studies on epidemiology, plant-pathogen interactions and development of disease mitigation methods such as breeding for resistance and crop rotation practices.

## Acknowledgements

This study was partially supported by Canadian Agricultural Partnership (no. 601322), the Saskatchewan Barley Development Commission, the Saskatchewan Wheat Development Commission, the Manitoba Crop Alliance, and the Western Grain Research Foundation and the Saskatchewan Ministry of Agriculture-Agriculture Development Fund.

---

**Fig. S1.**
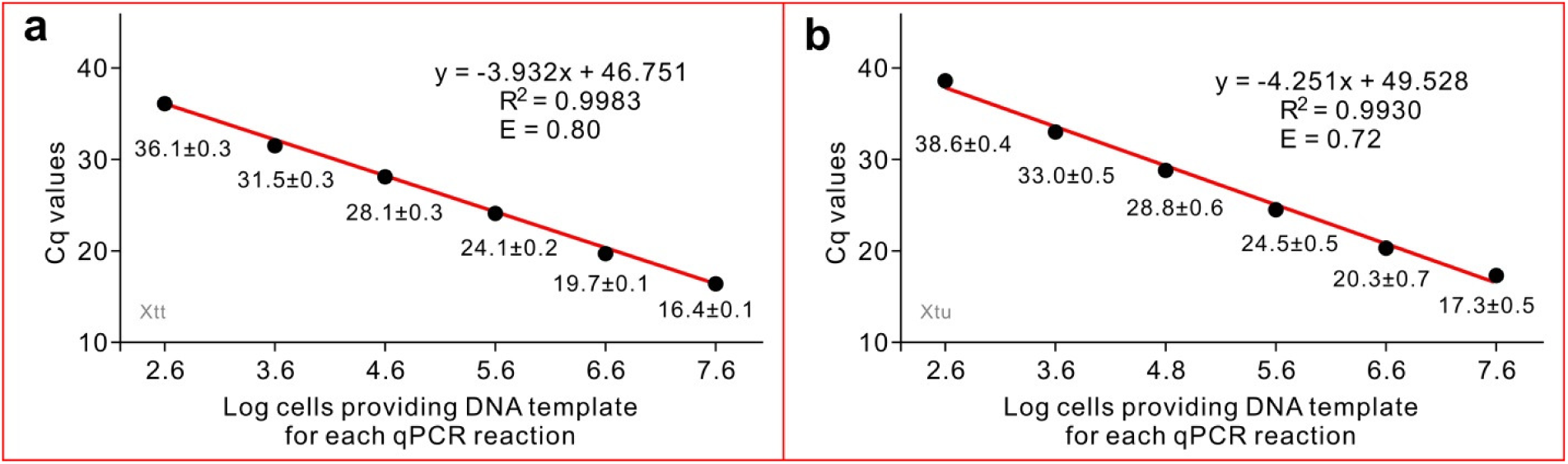
Sensitivity of the F11/R11/P11 qPCR system on the detection of *Xanthomonas translucens* pv. *translucens* (a) and pv. *undulosa* (b). This qPCR system was developed by Sarkes et al. (2022) and used in this study as a control to evaluate the sensitivity of the duplex qPCR system. The qPCR standard curves were generated from the mean of quantification cycle (Cq) values against log10 of the cell number from which the DNA was used in one reaction. The R^2^ score of the equation and the efficiency of the primers (E) are indicated over the curve. Efficiency was calculated as E = -1+10^(-1/slope)^. Each data point is shown as mean of three biological repeats ± standard deviation.

**Table S1.** *Xanthomonas translucens* strains isolated by the Alberta Plant Health Lab and used in this study

| Strain Name | Pathovar | Name in GenBank | GenBank accession number (length) |  |  |
| --- | --- | --- | --- | --- | --- |
|  |  |  | *cpn60* | 16S RNA | Whole genome<sup>a</sup> |
| CDCN-B045 | *undulosa* | CDCNb045 | MW759847 (609) | MW784564 (355) |  |
| CDCN-B046 | *undulosa* | CDCNb046 | MW759848 (616) | MW784565 (322) | JANVBQ010000026 |
| CDCN-B047 | *undulosa* | CDCNb047 | MW759849 (609) | MW784566 (338) |  |
| CDCN-B055 | *undulosa* | 1202-6-1-2 | OK663242 (556) |  |  |
| CDCN-B057 | *undulosa* | 1202-9-2-2 | OK663243 (556) |  |  |
| CDCN-B058 | *undulosa* | 1202-4-1-1 | OK663244 (556) |  |  |
| CDCN-B053 | *translucens* | 1219-B-1-3 | OK663240 (550) |  | JANVBP010000007 |
| CDCN-B054 | *translucens* | 1202-5-2-2 | OK663241 (550) |  |  |

<sup>a</sup>The whole genome sequencing was done by Dr. James Tambong, Agriculture and Agri-Food Canada, Ottawa Research and Development Centre.

**Table S2.**
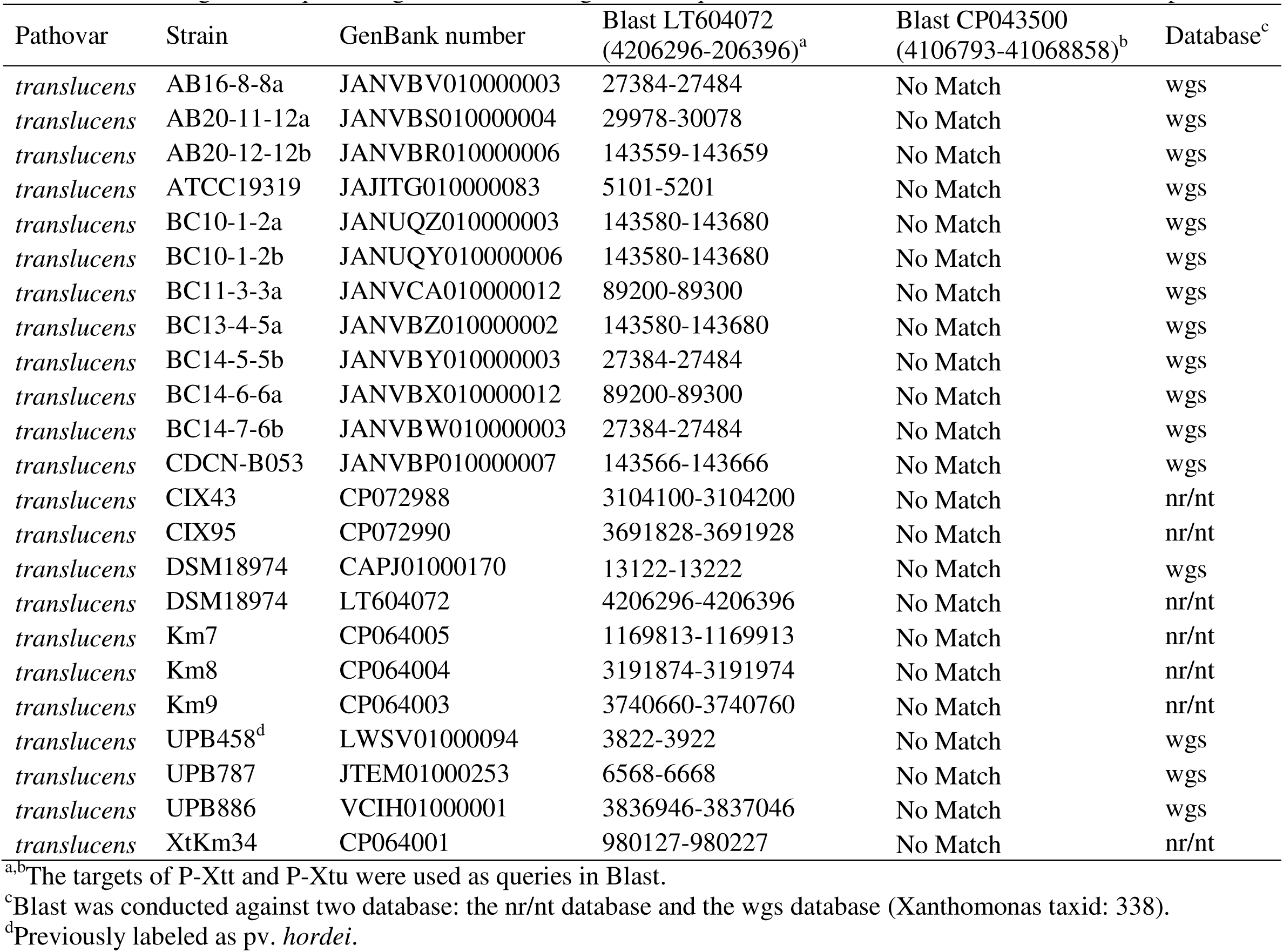
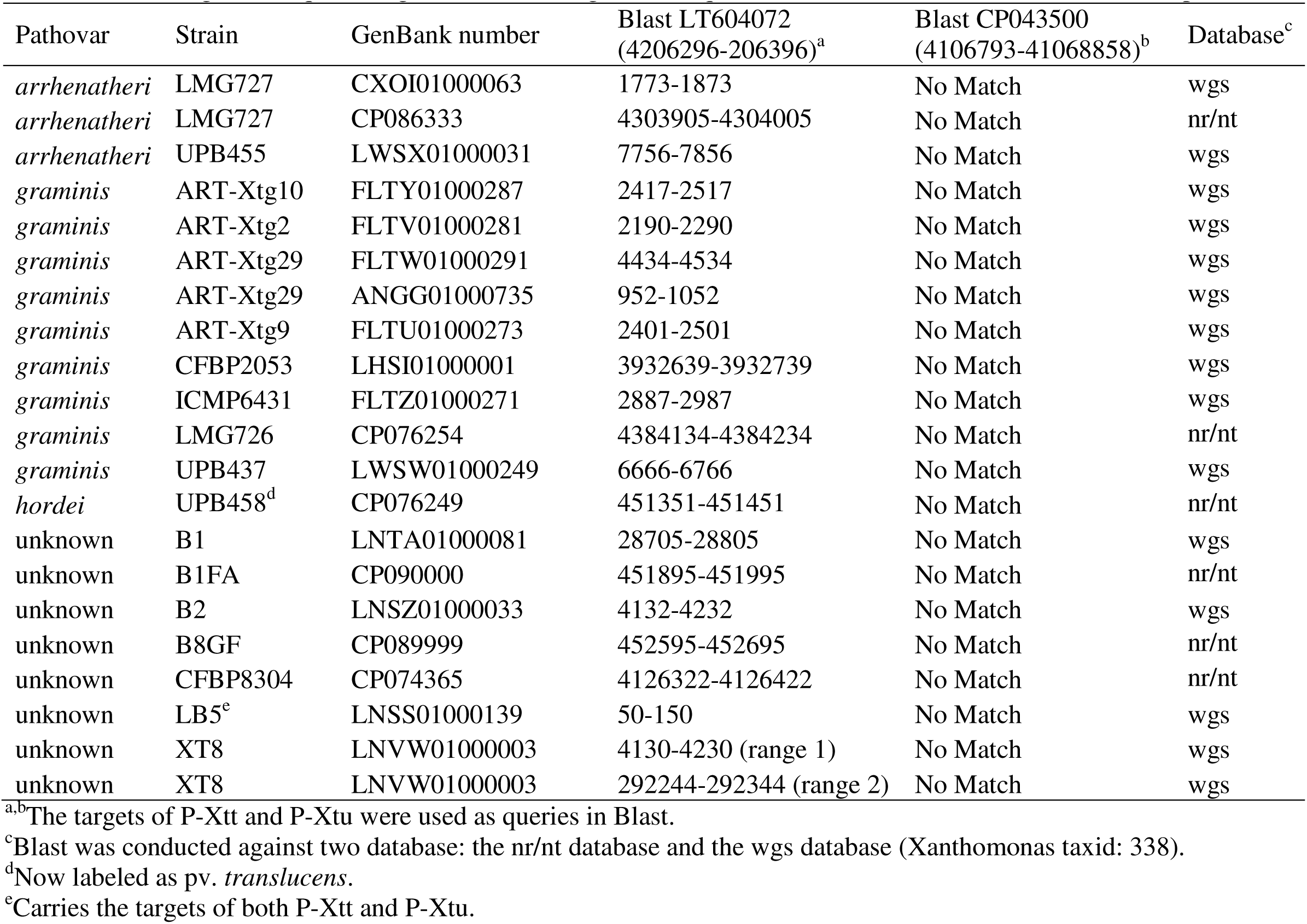
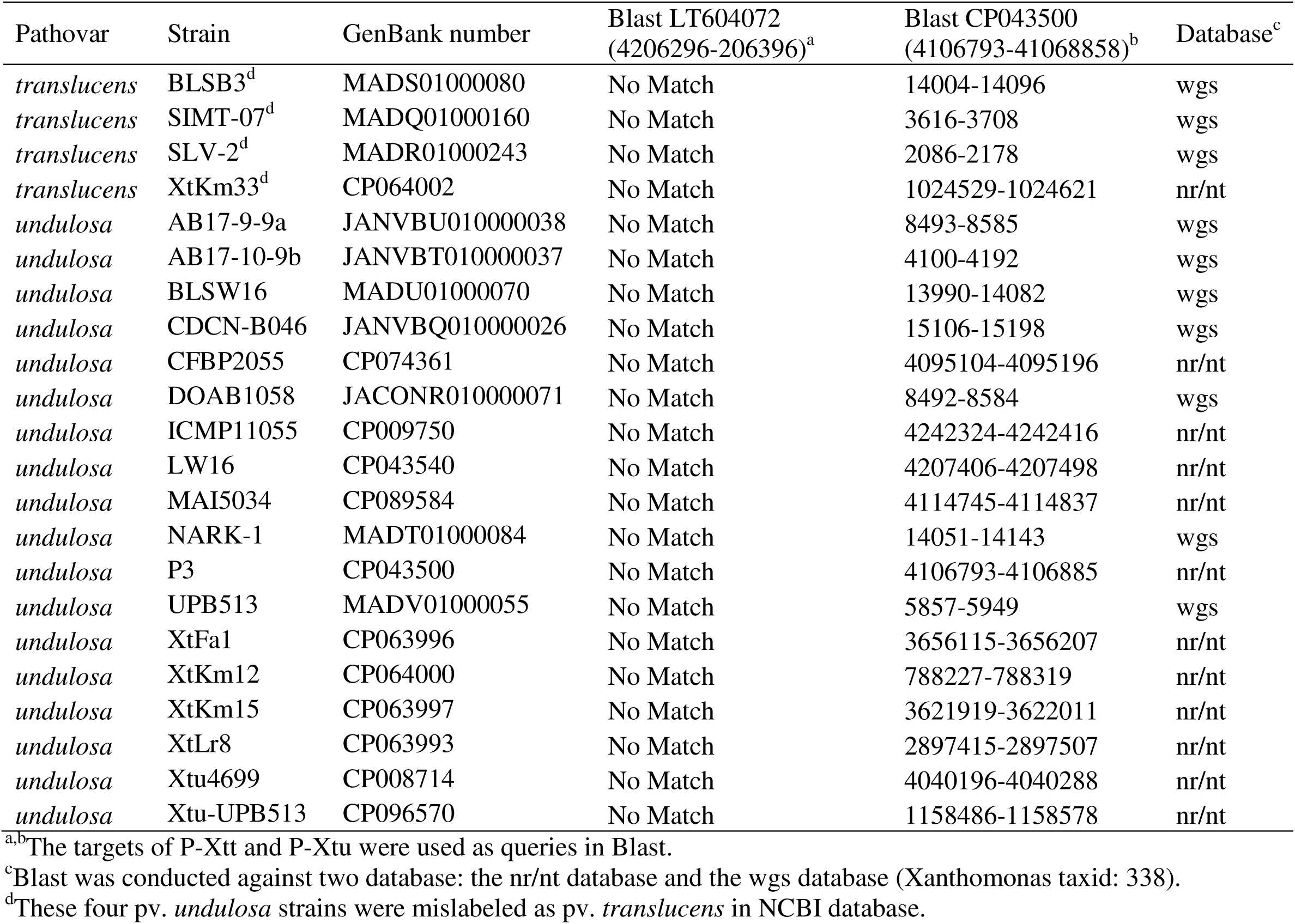
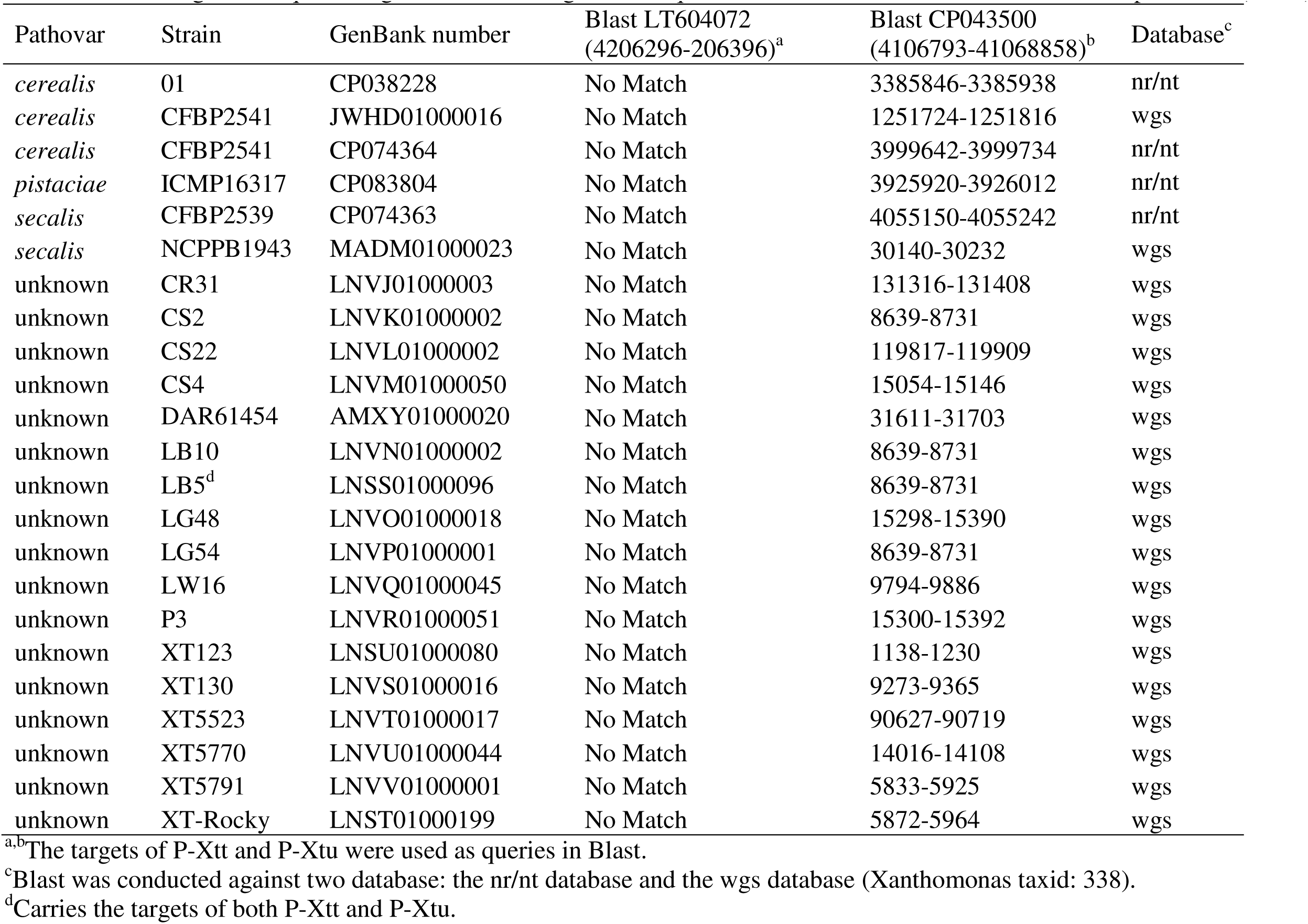
Orthologs of the qPCR targets in the whole-genome sequenced strains of *Xanthomonas translucens* pathovars

